# KlinkPPI: Single Point of Access to Protein-Protein Interactions Across Databases

**DOI:** 10.64898/2026.08.02.742057

**Authors:** Ahmad Lutfi, Sukrit Dang, Robert Warneke, Lutz Fischer, Juri Rappsilber

**Affiliations:** Technische Universität Berlin, Chair of Bioanalytics, 10623 Berlin, Germany; Si-M/”Der Simulierte Mensch”, a Science Framework of Technische Universität Berlin and Charité - Universitätsmedizin Berlin, Berlin, Germany; Indraprastha Institute of Information Technology Delhi, New Delhi, India - 110020

## Abstract

Protein–protein interaction (PPI) information is distributed across resources that differ in organism coverage, identifier systems, evidence models, confidence scores and access mechanisms, so assembling and comparing evidence for a protein requires source-specific queries, identifier conversion and extensive post-processing. We present KlinkPPI, a web server that retrieves, compares and exports PPI evidence from STRING, BioGRID, IntAct, CORUM, HuRI and Predictomes from a single query. KlinkPPI accepts UniProtKB accessions, NCBI Gene and Ensembl identifiers and gene names, and performs taxonomy-aware mapping to a common identifier space. Users can query individual proteins across all resources available for an organism, or retrieve organism-wide interaction sets. Results are presented per source so that database-specific evidence, annotations and confidence values are retained, while an integrated view exposes coverage and agreement between resources. KlinkPPI deliberately does not merge heterogeneous confidence scores, nor collapse functional associations, complex co-membership, binary interactions and structural predictions into a single consensus network. Results can be exported in PSI-MI TAB 2.8-compatible or Apache Parquet format with user-selected evidence fields. KlinkPPI is freely available at https://rappsilberlab.org/KlinkPPI/ and the source code at https://github.com/Rappsilber-Laboratory/KlinkPPI.

**Graphical abstract:** 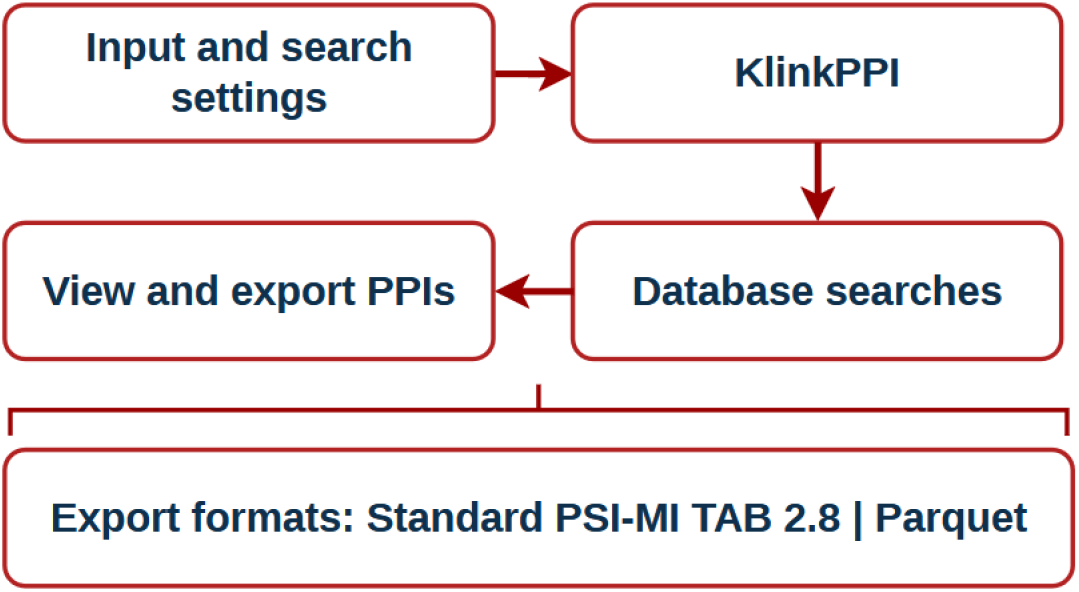

## Introduction

Protein–protein interactions (PPIs) form the molecular basis of many cellular processes, including signal transduction, protein-complex assembly, metabolism and the regulation of gene expression. PPI networks are therefore widely used to investigate protein function, characterize disease mechanisms and prioritize potential therapeutic targets^1,2^. Nevertheless, no single experimental method or database provides a complete representation of the interactome. Available PPI datasets differ substantially in organism coverage, experimental origin, biological context and the interpretation of what constitutes an interaction.

Reliable PPI datasets are also becoming increasingly important for machine-learning and deep-learning applications. Experimentally observed interactions can serve as training labels for interaction prediction, while PPI networks provide graph structures for representation learning and context-aware protein modelling. For example, D-SCRIPT was trained on more than 38 000 human PPIs to predict physical interactions directly from protein sequences and demonstrated cross-species generalization^3^. More recently, PLM-interact jointly encoded protein pairs and was trained using known human interaction data; it also used mutation-effect annotations from IntAct to predict whether sequence variants increase or decrease an interaction^4^. PINNACLE used context-specific PPI networks together with single-cell transcriptomic data to learn protein representations across tissues and cell types^5^. MINT extended protein language modelling to interacting protein sets by training on millions of curated physical interaction records derived from STRING^6^. These examples demonstrate that curated PPI information is not only a biological reference but also a fundamental training and benchmarking resource for modern artificial-intelligence models. Consequently, inconsistencies in identifiers, evidence definitions, duplicate records and dataset construction can propagate into model training and lead to biased or overly optimistic evaluations^7^.

Several established databases provide complementary views of protein relationships. STRING integrates experimentally supported and computationally inferred functional associations across many organisms^8^. BioGRID and IntAct provide curated molecular interactions derived from published experiments and submitted datasets^9,10^. CORUM focuses on manually curated mammalian protein complexes^11^, whereas the Human Reference Interactome, HuRI, provides a systematic map of experimentally detected human binary interactions^12^. Predictomes further expands the available interaction landscape with classifier-curated protein-interaction predictions generated using AlphaFold-based structural modelling^13^. However, these resources differ in their primary identifiers, evidence models, confidence measures, interaction definitions and output formats. Researchers may begin with UniProtKB accessions, NCBI Gene identifiers, Ensembl identifiers or gene names, while the queried database may use another identifier system. Although UniProtKB and Ensembl provide extensive cross-references^14,15^, querying and harmonizing multiple PPI resources still requires source-specific searches and substantial post-processing.

To address these challenges, we developed KlinkPPI, a web-based platform that provides unified access to STRING, BioGRID, IntAct, CORUM, HuRI and Predictomes. KlinkPPI accepts UniProtKB, NCBI Gene and Ensembl identifiers as well as gene names, performs taxonomy-aware identifier mapping and enables users to search one or several PPI resources through a single interface. Retrieved information is presented separately for each source, preserving database-specific evidence, annotations, confidence information and provenance. Users can inspect the returned interactions, select relevant output fields and export the results in the PSI-MI TAB 2.8-compatible tab-delimited^16^ format or in Parquet format for downstream analysis. KlinkPPI thereby reduces the manual effort required to retrieve and compare heterogeneous PPI information and supports the construction of transparent, reusable interaction datasets for biological analyses and data-driven modelling.

## Materials and methods

### PPI data sources

KlinkPPI integrates six complementary protein-protein interaction resources: STRING, BioGRID, IntAct, CORUM, the Human Reference Interactome (HuRI) and Predictomes. STRING provides experimentally supported and computationally inferred protein associations across a broad range of organisms. BioGRID and IntAct provide curated molecular interaction records derived from published experiments and submitted datasets. CORUM contains curated mammalian protein complexes, whereas HuRI provides a systematic map of experimentally detected human binary interactions. Predictomes contains classifier-curated protein-interaction predictions obtained from AlphaFold-Multimer-based structural screens.

Single-protein STRING queries were performed using the STRING version 12.0 REST API. IntAct interactions were retrieved through the EMBL–EBI IntAct web service. BioGRID records were obtained from the PSI-MI TAB file *BIOGRID-ALL-5*.*0*.*258*.*mitab*.*txt*. CORUM, HuRI and Predictomes were accessed through locally stored source files derived from their respective database releases. For complete-organism retrieval, KlinkPPI downloads and caches the STRING version 12.0 protein-links, protein-information and protein-alias files and the corresponding IntAct species-specific PSI-MI TAB archive where available.

The supported organisms were defined separately for each source using local taxonomy tables. These tables were combined into a unified species index containing the NCBI taxonomy identifier, a preferred display name, recognised species aliases and the databases available for that organism. Common names and abbreviated scientific names were included to support simple organism searches.

### Data retrieval and preprocessing

KlinkPPI supports two retrieval modes. In single-protein mode, users provide a protein or gene identifier, specify an organism or taxonomy identifier and select one or more interaction resources. The input is first resolved to a UniProtKB accession, after which a source-specific resolver retrieves and parses records from each selected database. Results from different sources are retained in separate database-specific result sets rather than being combined into a single consensus interaction list. In complete-organism mode, interaction datasets for the selected organism are processed as background jobs, with the processing state and number of retrieved pairs reported separately for each source.

Source-specific preprocessing was applied to preserve the semantics of each resource. STRING records were divided into interactions involving the query protein directly and additional edges between proteins returned in the query-centred network. The combined score and the neighbourhood, fusion, co-occurrence, experimental, co-expression, text-mining and database evidence channels were retained independently.

IntAct self-interactions were excluded from the single-protein results. Interactor pairs were ordered consistently and repeated records for the same pair were aggregated. The aggregated representation retains the number of supporting IntAct records, unique detection methods, PubMed identifiers, minimum and maximum feature counts and the IntAct MI score.

The BioGRID PSI-MI TAB file was preprocessed into an SQLite index to provide efficient accession-based retrieval. Only records containing UniProt/Swiss-Prot identifiers for both interactors were indexed. Each interaction was indexed in both orientations, allowing either protein to be used as the query. Detection method, interaction type, confidence value, gene name, taxonomy identifier and BioGRID record link were extracted from the original fields. The local index is rebuilt automatically when the source file or index schema changes.

HuRI interactions were loaded from a tab-separated interaction file and indexed by both Ensembl and, where available, UniProtKB identifiers. Records were indexed from both interaction orientations, and duplicate results were removed during retrieval. Predictomes records were indexed by UniProtKB accession, and interactions with a positive SPOC score were returned in descending score order. CORUM records were interpreted as protein-complex membership rather than evidence of direct binary binding. Associated complex subunits were reported together with the complex name, cell line and purification-method metadata.

### Identifier mapping and normalization

KlinkPPI accepts UniProtKB accessions, NCBI Gene identifiers, Ensembl identifiers and gene names. Non-UniProt identifiers are mapped to UniProtKB through the UniProt identifier-mapping API, with the supplied taxonomy identifier used to restrict the mapping where available. When the user does not specify an organism, the taxonomy identifier is obtained from the resolved UniProtKB entry. UniProtKB was selected as the principal query identifier because it provides extensive cross-references between protein, gene and organism-specific identifier systems.

Gene-name searches were performed against the UniProtKB search service using both the submitted gene name and the selected taxonomy identifier. Candidate entries were ranked by exact agreement with the reported gene symbols, followed by UniProt review status and accession. The user can therefore distinguish between multiple proteins associated with the same or similar gene names before performing the PPI search.

For HuRI queries, UniProtKB accessions were additionally mapped to Ensembl gene identifiers using the Ensembl cross-references provided in the corresponding UniProtKB entry. When no UniProtKB mapping was available for a HuRI interactor, the original Ensembl identifier was retained. This avoids discarding interactions merely because a cross-reference is unavailable. Ensembl provides complementary gene-centred annotation and identifier mapping across supported genomes.

Species names were normalized independently of protein identifiers. User-provided names were converted to a case-insensitive normalized representation and compared with scientific names, common names and database-specific aliases. Ambiguous species names produced candidate suggestions rather than being assigned automatically. The resolved NCBI taxonomy identifier was subsequently used to determine which PPI resources support the selected organism.

### Interaction normalization and evidence representation

Retrieved records were transformed into a common internal representation containing the two interactors, source database, organism information and source-specific evidence fields. For single-protein searches, the interaction partner is generally represented as interactor A and the query protein as interactor B. Original identifiers and source links are retained where available to allow users to inspect the corresponding database record.

KlinkPPI does not convert heterogeneous confidence values into a common numerical score. STRING evidence scores, IntAct MIscores, BioGRID confidence values and Predictomes scores have different definitions and are therefore retained in their original scales. Similarly, interactions are not merged across databases solely because they contain the same pair of identifiers. This design preserves source provenance and prevents distinct experimental, predicted and complex-membership evidence from being presented as equivalent observations.

For STRING, the combined association score and individual evidence channels are retained. IntAct records include detection methods, publication identifiers, feature counts, supporting-record counts and the MIscore. BioGRID records retain the experimental detection method, interaction type and confidence value. CORUM records retain complex-level annotations, including complex name, cell line and purification method. Predictomes records retain the SPOC score, KIRC score and number of unique structural contacts, while HuRI records preserve both UniProtKB and Ensembl identifiers where available.

Where an unambiguous mapping was available, source-specific interaction and detection terms were represented using PSI-MI controlled-vocabulary identifiers. Terms without a defined mapping were retained using their original source terminology. CORUM-derived pairs must be interpreted as co-membership of the same curated complex and not necessarily as evidence of a direct physical interaction between the two proteins.

### Data model and output formats

The backend returns a structured JSON response containing an input-information block and one result block for each selected database. Database-specific records remain separated throughout retrieval and presentation, enabling the web interface to display the available evidence, annotations and scores without removing source-specific fields. Users can export selected results as a tab-delimited molecular-interaction table or as Apache Parquet. The PSI-MI TAB export uses the standard interactor-identifier, taxonomy and source-database columns as its core fields: *#ID(s) interactor A, ID(s) interactor B, Taxid interactor A, Taxid interactor B, Source database(s)*.

In the Source database(s) field, KlinkPPI uses verified PSI-MI source-database terms where available: psi-mi:”MI:1014”(string), psi-mi:”MI:0463”(biogrid) and psi-mi:”MI:0469”(intact). CORUM, HuRI and Predictomes are reported using source names because no verified PSI-MI database accession was identified for these resources.

Optional fields are added according to the selected databases and evidence categories. These include STRING and Predictomes confidence values, IntAct publication identifiers and detection methods, BioGRID interaction types and experimental methods, and CORUM complex metadata. Multiple values within one field are separated using the pipe character. Missing values are represented by “-”, following the convention used in PSI-MI TAB files. PSI-MI TAB 2.8 provides a standardized tabular representation of molecular interactions, with one binary relationship represented per row.

Parquet exports retain source-specific columns in a columnar binary representation suitable for large-scale downstream analysis. Records are flattened into one row per reported interaction and include an explicit database field to preserve provenance. The files are generated using pandas and PyArrow.

### Database and software implementation

KlinkPPI uses a client-server architecture. The backend is implemented in Python using FastAPI and Pydantic for request handling and validation. Requests is used for communication with external APIs, pandas for tabular processing, SQLite for indexing the BioGRID source file and PyArrow for Parquet generation. The web interface is implemented using React and built with Vite. Tailwind CSS is used for responsive layout and visual styling. The interface supports source selection, single-protein and complete-organism search modes, organism suggestions, gene-name candidate selection, database-specific result views and configurable file export. The backend provides an endpoint for checking whether the KlinkPPI service is running, together with endpoints for species lookup, gene-name resolution, interaction retrieval, organism-level searches and data export. Complete-organism STRING and IntAct files are downloaded once and stored in a local cache for reuse. Frequently repeated taxonomy lookups, STRING links, Ensembl mappings and loaded STRING metadata are additionally cached in memory. The application can be installed and started using the project-level scripts, which create a Python virtual environment, install the backend and frontend dependencies and launch the FastAPI and Vite development servers.

## Database content and access

The overall workflow is summarized in Figure 1. The integrated resources contribute different types of protein relationships, ranging from curated experimental interactions and systematic binary interaction maps to protein-complex membership, functional associations and structurally predicted interactions. KlinkPPI preserves these distinctions instead of merging all retrieved records into an undifferentiated interaction set, allowing users to interpret and filter the results according to the requirements of their biological or computational application.

**Figure 1.**
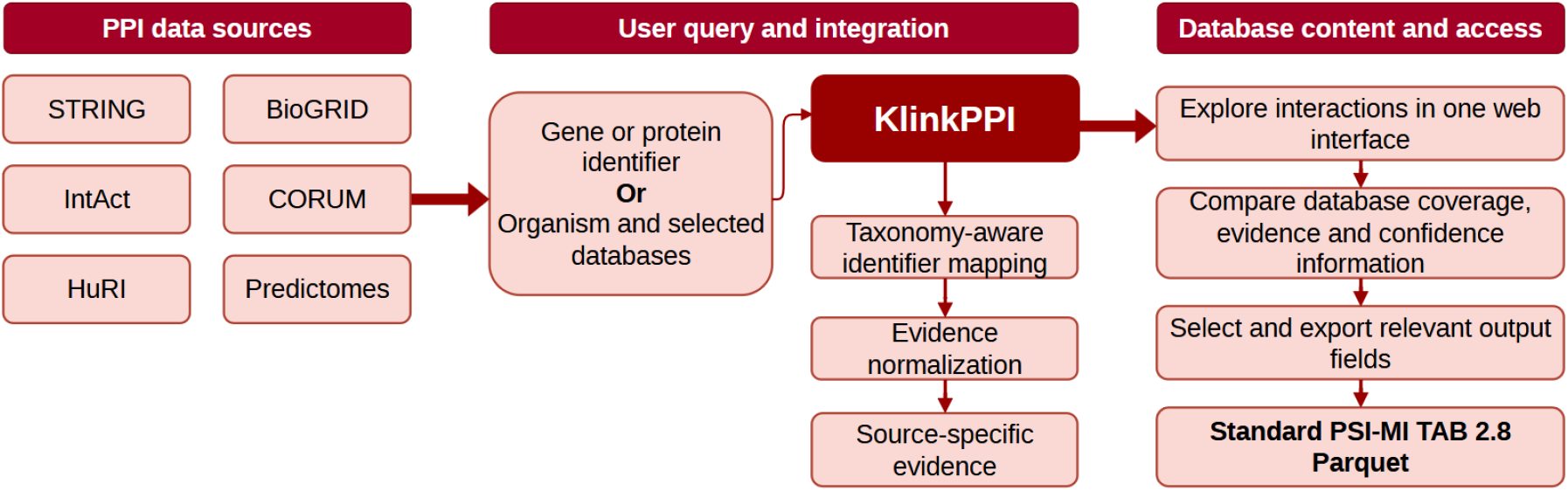
Overview of the KlinkPPI workflow and data-access architecture. KlinkPPI integrates PPI information from STRING, BioGRID, IntAct, CORUM, HuRI and Predictomes. Users can submit a gene or protein identifier or select an organism and supported databases for organism-wide retrieval. KlinkPPI performs taxonomy-aware identifier mapping and source-specific interaction processing while retaining evidence, confidence information and provenance. Retrieved interactions can be explored through the web interface and exported in the PSI-MI TAB 2.8 or Parquet format.

### Database content

KlinkPPI provides access to six PPI resources with markedly different taxonomic scopes (Figure 2). STRING offers the broadest coverage, supporting 12,536 NCBI taxonomy identifiers, followed by BioGRID with 211. IntAct and CORUM each support 16 taxonomy identifiers in the current implementation, whereas HuRI and Predictomes are restricted to human. The UpSet analysis further demonstrates that most taxonomy identifiers are exclusive to STRING, while a smaller core set is shared among multiple resources. These differences reflect the distinct aims of the integrated databases: STRING provides large-scale functional association networks across sequenced organisms, BioGRID and IntAct curate experimentally reported molecular interactions, CORUM focuses on mammalian protein complexes, HuRI provides a systematic human binary interactome and Predictomes contains human structurally predicted interactions. KlinkPPI uses this coverage information to display only databases available for the selected organism and to distinguish an unsupported resource from a supported resource returning no interactions.

**Figure 2.**
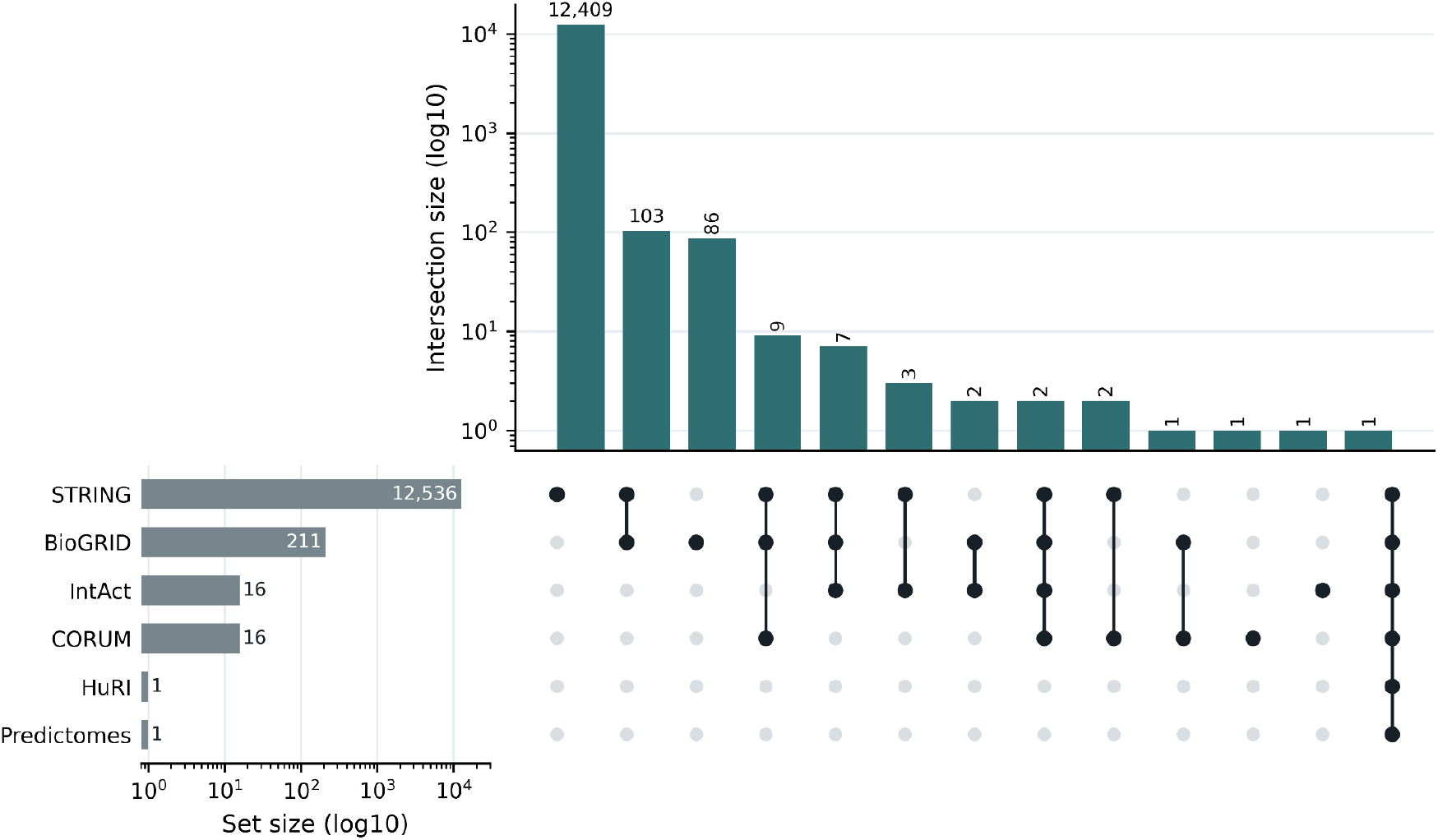
Taxonomic coverage of the PPI resources accessible through KlinkPPI. UpSet plot showing the overlap of supported NCBI taxonomy identifiers across STRING, BioGRID, IntAct, CORUM, HuRI and Predictomes. Horizontal bars indicate the total number of taxonomy identifiers supported by each resource, whereas vertical bars show the number associated with each resource intersection. Both set and intersection sizes are displayed on logarithmic scales. STRING provides the broadest taxonomic coverage, supporting 12,536 taxonomy identifiers, followed by BioGRID with 211. IntAct and CORUM each support 16 taxonomy identifiers, whereas HuRI and Predictomes are restricted to human in the current KlinkPPI release.

### Web interface

The KlinkPPI web interface guides users through input specification, identifier resolution, database selection, result inspection and data export. Users can initiate a protein-centred search using a UniProtKB accession, NCBI Gene identifier, Ensembl identifier or gene name. A species name or NCBI taxonomy identifier can be supplied to restrict identifier mapping and prevent assignments to proteins from an unintended organism. For gene-name queries, matching UniProtKB candidates are presented to the user before the interaction search is initiated. Users may query one database or several resources simultaneously. After processing, an input summary reports the normalized UniProtKB accession, selected organism and queried databases. Results are displayed in separate, source-specific panels for STRING, BioGRID, IntAct, CORUM, HuRI and Predictomes. The interface distinguishes between successful retrieval, absence of interactions and lack of organism support, thereby preventing an unsupported database from being interpreted as evidence that no interaction exists. Each result panel displays fields relevant to the corresponding resource. These may include interactor identifiers, gene names, interaction or detection methods, publication identifiers, confidence values, complex information and links to the original database records. Keeping the sources in separate panels enables users to inspect both agreement and disagreement between databases without losing the original evidence context.

A representative protein-centred query is shown in Figure 3. The human gene VAC14 was submitted together with the organism Homo sapiens, specified either by name or by the NCBI taxonomy identifier 9606, and all six interaction resources were selected. Before retrieving interaction data, KlinkPPI resolved the gene-name query to the corresponding UniProtKB accession, Q08AM6. The results were then displayed in separate, database-specific panels. The Shared Interaction Log provides an overview of the retrieved records, including the total number of interactions, the numbers of shared and resource-specific pairs, and the overlap patterns among the selected databases (Figure 3A). The source-specific panels report the interactions or associations involving VAC14 together with the evidence fields and scoring metrics provided by each resource (Figure 3B). Because the databases use different interaction definitions, evidence types and scoring systems, their outputs are not merged into a single confidence score. Instead, the separate presentation enables users to compare the supporting evidence across resources and assess the consistency and provenance of individual interactions.

**Figure 3.**
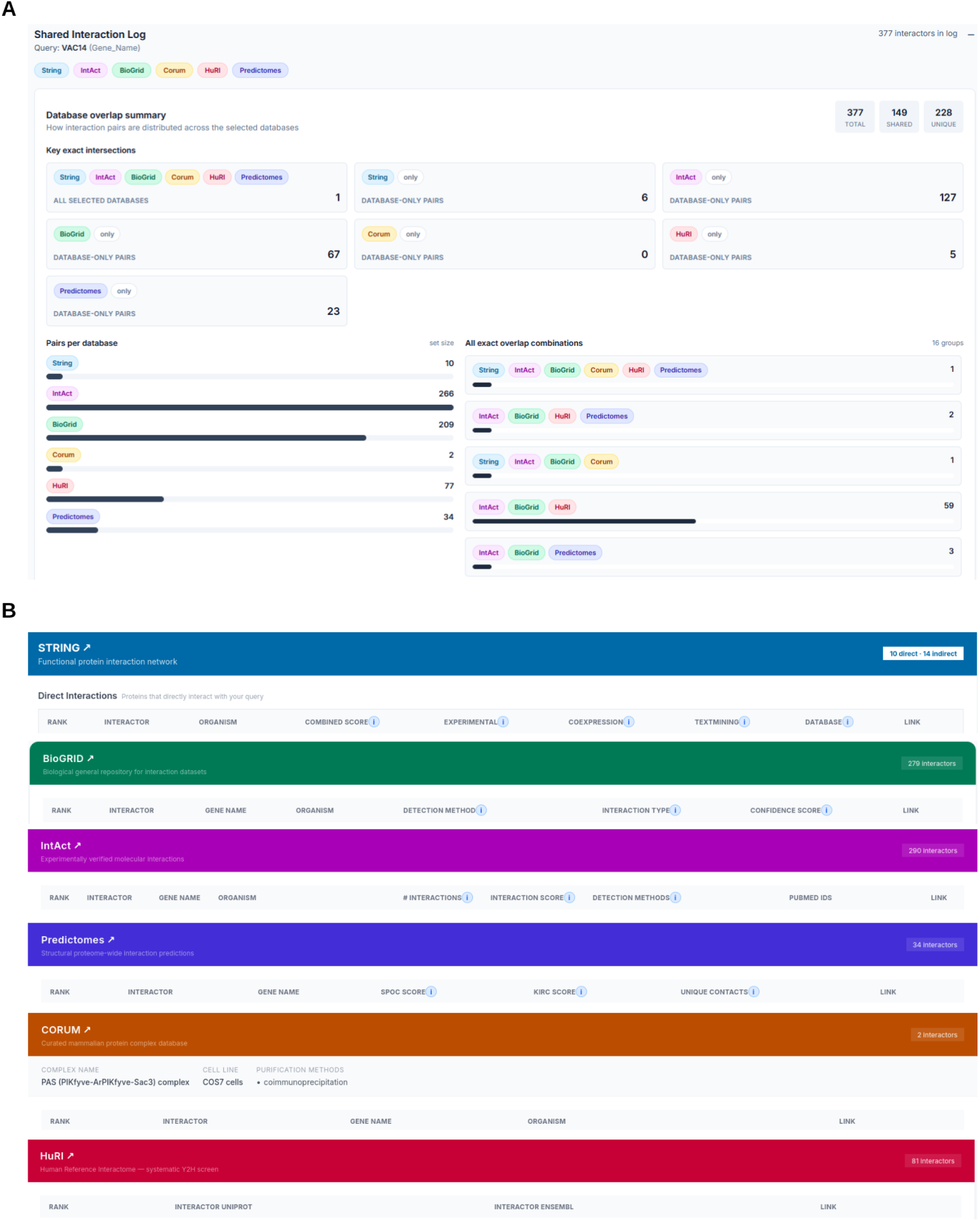
Cross-database interaction overview and source-specific presentation of VAC14 interaction results in KlinkPPI. **(A)** Shared Interaction Log for the human gene VAC14, integrating results from STRING, IntAct, BioGRID, CORUM, HuRI and Predictomes. The overview summarizes 377 retrieved interaction records, comprising 149 interactions shared between at least two resources and 228 resource-specific interactions. It also reports the number of pairs contributed by each database, highlights interactions found in all selected resources or exclusively in a single resource, and displays the observed combinations of database overlap. **(B)** Source-specific result panels retain the structure and evidence fields provided by each resource. The example shows 10 direct and 14 indirect STRING associations, 279 BioGRID interactors, 290 IntAct interactors, 34 Predictomes predictions, two CORUM complex-derived interactions and 81 HuRI interactors. Results are presented with database-appropriate metadata, including interaction scores, evidence channels, detection methods, publication identifiers, complex information and links to the corresponding source records.

### Use cases

A primary use case of KlinkPPI is the investigation of the evidence supporting a protein or protein pair across multiple resources. A user can submit a protein identifier, select all databases available for the corresponding organism and inspect the retrieved results in parallel. An interaction may, for example, be supported by curated experimental records in BioGRID and IntAct, appear as a systematic binary interaction in HuRI, occur between members of a CORUM complex and receive additional support from STRING or Predictomes.

KlinkPPI retains the evidence associated with each observation. The user can therefore examine the reported experimental methods and interaction types in BioGRID, the number of IntAct records and supporting publications, the individual STRING evidence channels and the structural prediction scores provided by Predictomes. This allows interactions supported by several independent sources to be distinguished from relationships reported by only one database. At the same time, the separate presentation prevents complex co-membership or functional association from being mistaken automatically for evidence of direct physical binding.

This use case is relevant when selecting candidate interactions for experimental validation. Candidates may be prioritized according to the type and amount of existing evidence; for example, by selecting interactions supported by several publications, interactions with experimental support in STRING and IntAct, or structurally plausible predictions with high SPOC or KIRC scores and multiple predicted contacts. KlinkPPI does not impose one ranking rule; instead, it supplies the evidence required for the user to define a transparent prioritization strategy.

KlinkPPI can be used to construct provenance-aware datasets for machine-learning and deep-learning models of PPIs. Protein language models and sequence-, structure- or graph-based interaction predictors require carefully defined positive interaction examples. Through complete-organism retrieval and Parquet export, users can collect source-specific interaction records while retaining identifiers, organisms, evidence types and confidence information. Different positive sets can be created according to the modelling objective. A model intended to predict direct binary binding could be trained primarily on HuRI interactions and experimentally supported records from IntAct or BioGRID. CORUM co-membership could instead be used for learning protein-complex organization, while STRING associations could support models of broader functional relationships. Predictomes records can provide structurally supported candidate interactions or an independent score for comparison with sequence-based predictions. Keeping these categories separate reduces the risk of training a model on biologically heterogeneous labels. The reported scores can also be incorporated into the learning strategy (weights). High-confidence records can be selected using source-specific thresholds, or confidence values can be used as sample weights rather than reducing all interactions to equally weighted binary labels. Individual STRING evidence channels can be supplied as separate features, allowing a model to distinguish experimental, genomic-context, co-expression and text-mining support. IntAct record counts, methods and publication identifiers can describe the multiplicity and diversity of experimental evidence. Predictomes scores and contact counts can provide structural features for multimodal models.

A further use case is the construction of interaction networks in which edges retain their original biological interpretation. Users can retrieve all available interactions for an organism and construct separate or multilayer networks for binary interactions, curated experimental interactions, functional associations, complex co-membership and structural predictions. The database field and source-specific annotations allow these edge classes to be analysed independently or combined using explicitly defined rules.

Such networks can be used for graph-based machine learning, protein-function prediction, candidate-gene prioritization or community detection. Confidence scores can be used as source-specific edge weights, while experimental methods, publication support or structural contacts can be represented as edge attributes. A graph neural network could, for example, treat STRING evidence channels as multidimensional edge features rather than reducing them to one combined value. Alternatively, separate graph layers could represent IntAct or BioGRID experimental interactions, CORUM complex relationships and Predictomes structural predictions.

## Results

KlinkPPI successfully integrated protein-interaction retrieval from six resources within a common query and presentation workflow. The integrated databases showed markedly different taxonomic coverage, ranging from 12,536 NCBI taxonomy identifiers in STRING to human-specific coverage in HuRI and Predictomes (Figure 2). This coverage information was used during query configuration to identify which resources were available for the selected organism and to distinguish an unsupported database from a supported database returning no interactions. The representative VAC14 query demonstrated the complete protein-centred workflow (Figure 3). A gene-name query was first resolved to UniProtKB accession Q08AM6 using the specified human taxonomy. KlinkPPI then queried the selected resources and presented their results separately. STRING returned 10 direct and 14 indirect associations. The direct associations were not represented solely by a combined confidence score; the individual evidence channels were retained, revealing substantial differences in the types of support underlying similarly ranked interactions. For example, some associations received strong experimental or curated-database support, whereas others were predominantly supported by text mining or co-expression. The interface therefore exposes evidence composition as well as overall ranking.

Equivalent source-specific representations were generated for the other integrated resources. BioGRID results retained experimental methods and interaction types, whereas IntAct results included detection methods, supporting publications, record counts and MI scores. CORUM results described shared membership of curated complexes, HuRI contributed systematically detected binary human interactions and Predictomes reported structural prediction scores and predicted contact information. Keeping these outputs separate prevented fundamentally different relationship classes from being interpreted as equivalent evidence.

Retrieved records could be exported with user-selected fields in PSI-MI TAB 2.8 or Parquet format. Database provenance and original score names were preserved in both formats. This enabled the exported data to be filtered according to the intended downstream application, including experimental candidate prioritization, interaction-network construction and the preparation of training or evaluation datasets for machine-learning.

## Discussion

KlinkPPI addresses a recurring practical challenge in PPI research: relevant interaction information is distributed across resources that differ in identifiers, organisms, evidence models, confidence scores and access mechanisms. The principal contribution of KlinkPPI is therefore not merely the aggregation of protein pairs, but the provision of a common access workflow while preserving the biological interpretation and provenance of each source. This is important because a functional association in STRING, a curated experimental record in IntAct or BioGRID, a binary interaction in HuRI, a shared CORUM complex and a structural prediction in Predictomes do not represent equivalent observations.

The retention of source-specific evidence is particularly valuable for machine-learning applications. PPI prediction models are sensitive to the construction of positive examples, the definition of negative examples and information leakage between training and test sets. KlinkPPI enables users to select interaction classes that match the intended modelling objective. Binary or experimentally detected interactions can be separated from functional associations and complex co-memberships, while source-specific confidence scores can be used for filtering, sample weighting or auxiliary features. STRING evidence channels can be represented as multidimensional edge attributes, IntAct publication and detection-method information can describe evidence multiplicity, and Predictomes scores and contact counts can provide structural information for multimodal models. KlinkPPI deliberately avoids combining these values into a single global score because they are not calibrated measurements of the same quantity.

Future development will focus on automated and versioned database updates with the integration of more databases, and including a documented programmatic API, optional Python and R clients, and tools for comparing cross-database agreement.

## Code availability

The source code for KlinkPPI is openly available at https://github.com/Rappsilber-Laboratory/KlinkPPI. The repository contains the backend and frontend source code, dependency specifications and instructions for local installation and execution.

## Data availability

The release source code with all related databases are in Zenodo. https://doi.org/10.5281/zenodo.21647179.

## Funding

This work was supported by the Deutsche Forschungsgemeinschaft (DFG, German Research Foundation) under Germany’s Excellence Strategy - EXC 2008 - 390540038 - UniSysCat and the European Research Council under the European Union’s Horizon Europe research and innovation programme (grant agreement No. 101119142 - TransFORM).

## Author contributions

A.L. implemented the new version of the KlinkPPI and prepared the initial draft. S.D. built the first tool prototype. R.W. and L.F. contributed to tool testing and manuscript revision. J.R. conceived and supervised the project and revised the manuscript.

## Competing interests

The authors declare no competing interests.

## Notes

https://doi.org/10.5281/zenodo.21647179.

### Competing Interest Statement

The authors have declared no competing interest.

